# Supercharging enables organized assembly of synthetic biomolecules

**DOI:** 10.1101/323261

**Authors:** Anna J. Simon, Vyas Ramasubramani, Jens Glaser, Arti Pothukuchy, Jillian Gerberich, Janelle Leggere, Barrett R. Morrow, Jimmy Golihar, Cheulhee Jung, Sharon C. Glotzer, David W. Taylor, Andrew D. Ellington

## Abstract

There are few methods for the assembly of defined protein oligomers and higher order structures that could serve as novel biomaterials. Using fluorescent proteins as a model system, we have engineered novel oligomerization states by combining oppositely supercharged variants. A well-defined, highly symmetrical 16-mer (two stacked, circular octamers) can be formed from alternating charged proteins; higher order structures then form in a hierarchical fashion from this discrete protomer. During SUpercharged PRotein Assembly (SuPrA), electrostatic attraction between oppositely charged variants drives interaction, while shape and patchy physicochemical interactions lead to spatial organization along specific interfaces, ultimately resulting in protein assemblies never before seen in nature.

---

The assembly of genetically-encoded molecules into symmetric, supramolecular materials enables complex functions in natural biosystems and would likewise prove valuable in the development of bionanotechnologies including drug delivery, energy transport, and biological information storage.^1–29^ alone can enable formation of complex symmetrical assemblies. Shape complementarity has likewise been exploited to arrange synthetic linear biomolecules, i.e., double-stranded DNA strands, into distinct structures, demonstrating its utility for for the higher order arrangement of synthetic linear biomolecules, suggesting a utility beyond inorganic systems^30,31^.

Herein we demonstrate a simple, robust strategy to assemble normally monomeric proteins into well defined, oligomeric quaternary structures and micron-scale particles. This strategy centers on driving protein interactions by engineering oppositely supercharged variants. Generally, oppositely-charged proteins interact through simple electrostativ interactions.^37^ and Matryoshka-like cages^38^ from naturally assembling proteins. Here, we suspected that shape and physicochemical features would favor assembly of oppositely charged protein pairs along particular interfaces to produce defined oligomeric assemblies (**Fig. 1a**). To the extent that these hypotheses can be realized, the ability to readily engineer synthetic, scalable molecular assemblies into almost any protein via supercharging should prove useful for technologies ranging from pharmaceutical targeting to artifical energy harvesting to “smart” sensing and building materials^1, 16, 17, 39^.

**Fig. 1.**
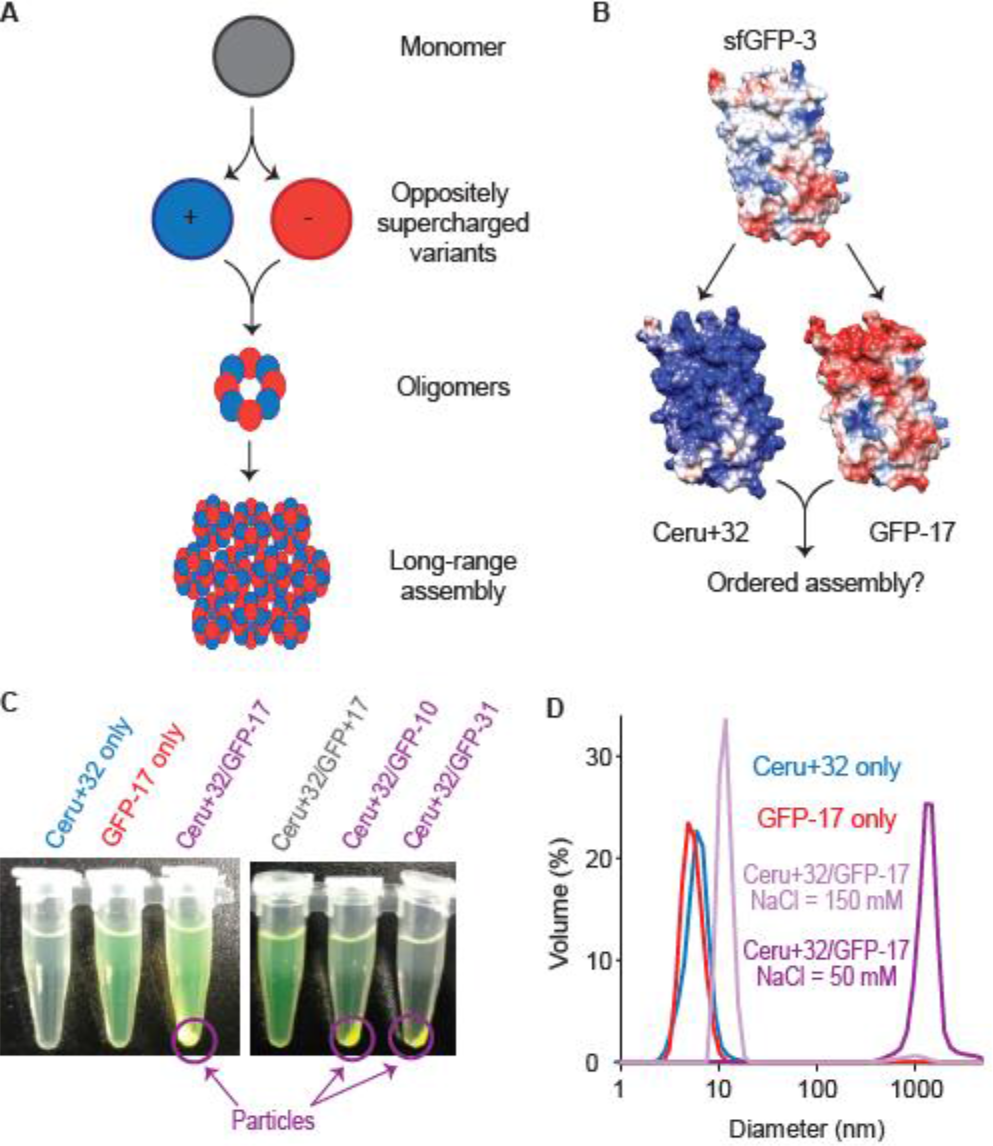
Oppositely supercharged Cerulean and GFP variant proteins as a model system for charge-mediated protein assembly. **a,** We hypothesized that generating and mixing oppositely supercharged variants of normally monomeric proteins could produce symmetrical oligomeric and higher-order structures **b,** To investigate the use of supercharging for the ordered assembly of proteins we employed supercharged GFP variants engineered by mutating surface residues of superfolder GFP to positively or negatively charged residues, shaded blue and red, respectively. For the negative protein we employed a cerulean variant of superfolder GFP+3, “Ceru+32”;^40^ for the negative variants we employed Rosetta Supercharged GFPs with negative charges of −10, −17, and −31.^41^ **c,** While all of the individual proteins are easily soluble and do not produce precipitant, mixing 0.1 mg/ml Ceru+32 with an equivalent amount any of the negatively supercharged GFPs at 50 mM produces visible particles that sink upon light centrifugation. Mixing Ceru+32 with a positively supercharged control protein, GFP+17^41^ does not produce precipitation. **d,** Dynamic Light Scattering (DLS) measurements indicate that the precipants formed from mixing Ceru+32 and GFP-17 at 50 mM NaCl are largely composed of monodisperse particles with an average diameter of 1421±136 nm. Mixing Ceru+32 and GFP-17 at 150 mM NaCl, in contrast, produces monodisperse particles with an average diameter of 11.6±1.8 nm. Measurements of Ceru+32 and GFP-17 alone indicate particles of average diameter of 6.3±1.6 nm and 5.5±1.4 nm, consistent with monomers. Measurements of Ceru+32/GFP-10 and Ceru+32/GFP-31 and GFP-10 and GFP-31 alone indicate slightly larger and more polydisperse mixed particles.

As a model system we employed variant pairs of monomeric supercharged green fluorescent proteins (GFPs) containing excess basic or acidic amino acids on their surfaces (**Fig. 1b**, **Supplementary Fig. 1**).^47^ ultimately yielding “Ceru+32.” Native gels (**Supplementary Fig. 2**) and absorbance/emission spectra (**Supplementary Fig. 3**) indicated that each protein expressed monomerically and with expected fluorescent properties. Solutions containing individual proteins showed no precipitation (**Fig. 1c**), and Dynamic Light Scattering (DLS) measurements indicated diameters of sizes of 5.7±1.7, 6.2±1.9, 5.5±1.4, and 5.9±1.9 nm for Ceru+32, GFP-10, GFP-17, and GFP-31, consistent with the size of GFP monomers (**Fig 1d**, **Supplementary Fig. 4**).

While each GFP variant was individually monomeric, combining oppositely-charged proteins produces larger particles. Mixing equal amounts of Ceru+32 and any of the negatively supercharged GFPs at 50 mM NaCl produced a cloudy solution containing fluorescent particles that settled after brief centrifugation at 1700 x g relative centrifugal force (**Fig. 1c**). Dynamic Light Scattering (DLS) measurements indicated that Ceru+32/GFP-10 and Ceru+32/GFP-17 were micron-scale and quite monodisperse, with average diameters of 1675±251 nm and 1421±136 nm, respectively (**Fig. 1d**, **Supplementary Fig. 4**). Ceru+32/GFP-31 particles were larger and more polydisperse, with an average diameter of 2452±984 nm (**Supplementary Fig. 4**). Mixing two positively supercharged proteins (Ceru+32/GFP+17) produced no visible precipitation (**Fig. 1c**).

The observed interaction of opposite-but not like-charged proteins indicates that assembly is mediated by electrostatic interactions, and we therefore next investigated the relationship between particle size and ionic strength. Notably, combining Ceru+32 and GFP-17 at higher ionic strength yields smaller but definitively non-monomeric particles, producing, for example, a particle size of 11.6±1.8 nm at 150 mM (**Fig. 1d**). Measuring Ceru+32/GFP-17 particle size over a range of NaCl concentrations indicates a phase-transition behavior, with consistent diameters of ~1400 nm at NaCl ≤50 mM and ~12 nm at 100-300 mM NaCl (**Fig. 1d**, **2a**). Ceru+32/GFP-17 particle diameter decreases slightly to ~9 nm at NaCl > 300 mM, suggesting that at these high-salt conditions a fraction of the ~12 nm particle may dissociate. Formation of the 11 nm-scale protomers from the larger particles is reversible. Addition of high-salt buffer to increase sodium chloride concentration from 50 mM to 100 mM reduced the average diameter of Ceru+32/GFP-17 particles from 1280±327 nm to 12.5±2.2 nm (**Fig. 2b**). This transition occurs rapidly, with the solutions becoming clear and particle size decreasing immediately upon addition of salt. This reversibility suggests that the micron scale particles assemble via well-defined, discrete interactions between folded structures, as opposed to non-specifically via unfolded, irreversible interactions.^48^ Ceru+32/GFP-10 and Ceru+32/GFP-31 particles show qualitatively similar salt-dependence as Ceru+32/GFP-17 (**Supplementary Fig. 5**). However, compared to Ceru+32/GFP-17 the size transition from ~1300 nm to ~12 nm diameter occurs at considerably higher NaCl concentrations, 150 mM and 300 mM for GFP-10 and −31, respectively, indicating that the larger particles may have even higher stabilities.

**Fig. 2.**
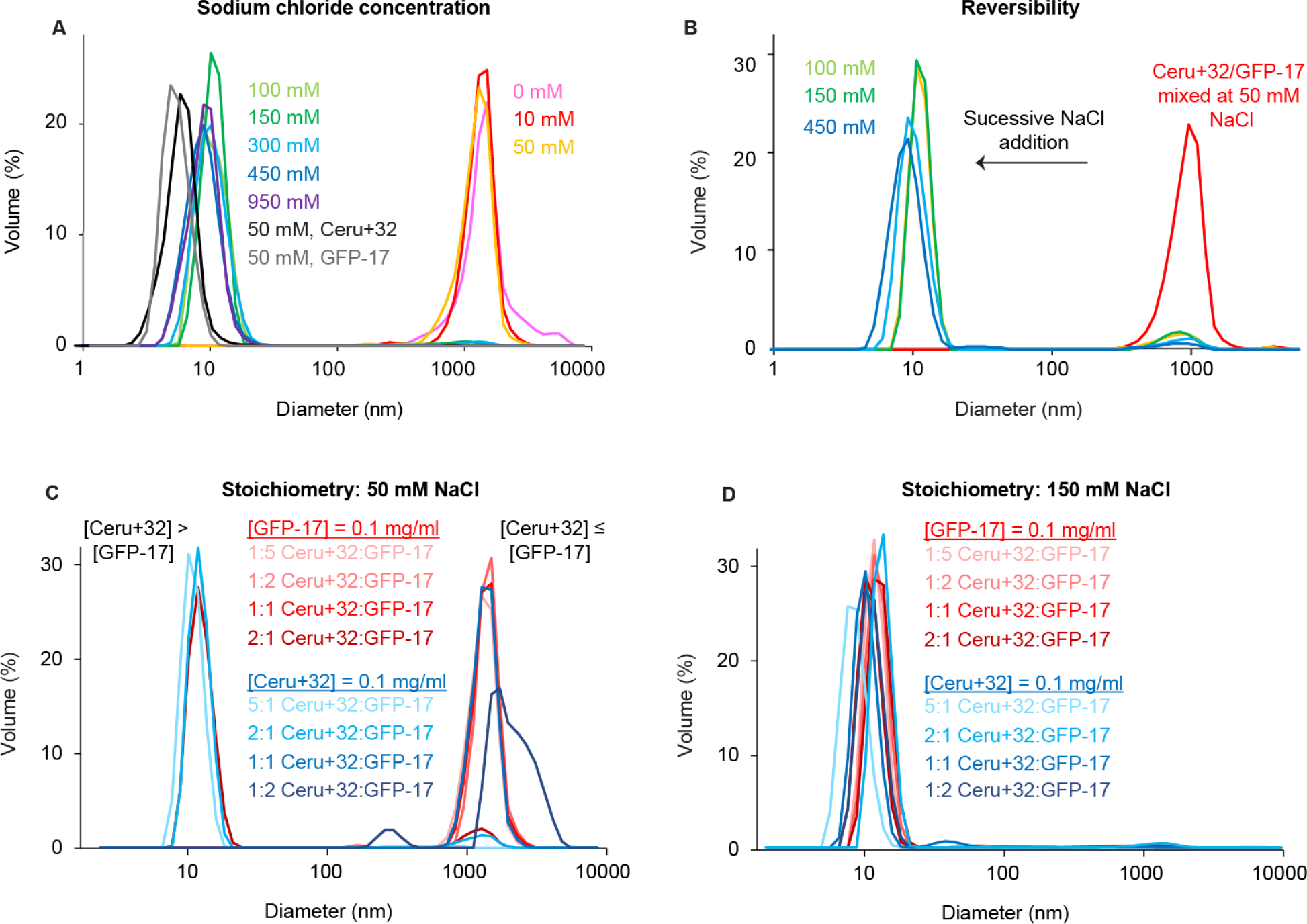
Ceru+32/GFP-17 particle size depends on environmental conditions. **a,** Particle size depends on sodium chloride in a phase-like manner. At pH 7.4 and 0.1 mg/ml of both Ceru+32 and GFP-17, ~1300 nm scale particles assemble when mixed at sodium chloride concentrations at and below 50 mM; ~12 nm particles assemble when mixed at sodium chloride concentrations at and above 100 mM. Even at the highest sodium chloride concentration we measure, 950 mM, particle size remains significantly higher than that of the monomeric size measured in solutions containing single proteins. **b,** Micron scale particle formation was reversible: increasing the sodium chloride concentration of Ceru+32/GFP-17 particles initially mixed at 50 mM to 100 mM rapidly reduced particle size from 1280 ± 327 nm to 12.5 nm ± 2.2 nm. **c,** Micron vs ~12 nm particle formation depends on stoichiometry: at 50 mM NaCl with excess Ceru+32, ~12 nm particles form; with equivalent or excess GFP-17, ~1300 nm particles form. **d,** At 150 mM NaCl, at 5:1 ratios of Ceru+32 to GFP-17, ~7 nm particles form; at all other measured ratios, ~10-12 nm particles form.

Ceru+32/GFP-17 particle size likewise depends on protein stoichiometry and pH. To characterize the effects of stoichiometry, we measured the size of particles formed at 50 mM NaCl and 150 mM NaCl from 0.1 mg/ml of either Ceru+32 or GFP-17 and 0.02, 0.05, 0.01, or 0.2 mg/ml of the other. At 50 mM NaCl, DLS measurements indicated ~1300 nm particles at excess or equivalent GFP-17 but ~10-12 nm particles at excess Ceru+32 (**Fig. 2c**). At 150 mM NaCl, DLS measurements indicate ~12 nm particles at 1:2, 1:1, 2:1, and 5:1 ratios of Ceru+32 to GFP-17 and ~7 nm particles at 1:5 ratios of Ceru+32 to GFP-17. Although the DLS measurements did not generally directly indicate the presence of monomeric proteins even at unequal stoichiometries, we suspect that they are likely present: DLS generally provides the “major” component in a solution via extrapolating particles sizes from the time-correlation of scattering patterns. Because scattering intensity depends on particle diameter to the 6^th^ power, DLS contains a significant bias toward larger components,^49^ suggesting that our measurements may resolve only the largest major component (i.e., the 12 nm particle), or potentially, a size in between two similarly sized but different components (i.e., the ~6 nm protein and ~12 nm particle). Decreasing pH dissolved the both the protomer and larger particle completely to monomers. At 50 mM NaCl, particle size decreased from 1622±556 nm at pH 7.4 to 5.7±2.4 nm at pH 6.5; at 150 mM NaCl, particle size decreased from 11.0±3.1 nm at pH 7.4 to 6.0±0.7 nm (**Supplementary Fig. 6**). Fluorescence measurements confirmed that both proteins remain well-folded at pH 6.8, suggesting that dissolution occurs due to decreased interaction favorability rather than unfolding (**Supplementary Fig. 7**). Taken together, these observations that mixtures of Ceru+32 and GFP-17 assemble into three distinct phases ~6 nm (monomeric), ~12 nm, and ~1300 nm - at a broad range of NaCl, stoichiometric, and pH conditions suggest that the proteins assemble via discrete, specific intermolecular interactions.

To study the architecture of the ~12 nm particles we visualized negatively stained Ceru+32/GFP-17 particles formed at 150 mM NaCl using electron microscopy. Raw electron micrographs show mono-disperse, globular protomers with dimensions of ~160-Å in diameter that appear to contain internal features (**Supplementary Fig. 8**). Intriguingly, the particles show 8-fold symmetry after reference-free alignment, classification, and averaging to increase the signal-to-noise ratio (**Supplementary Fig. 8**). Using 3D reconstruction techniques (see **Materials and Methods**) we obtained a structure of this complex at ~18-Å resolution clearly showing eight globular densities of the same size arranged into a single ring (**Fig. 3a**). There appear to be two symmetric rings stacked on top of one another and offset by about half a subunit, ultimately forming a barrel (**Fig. 3b**). Each globular density in the particles in the rings corresponded to the size and shape of a GFP monomer; the x-ray crystal structure of GFP can be easily accommodated within the density (**Fig. 3b**). Not This structure is roughly equivalent in volume to the ~11 nm diameter sphere measured by DLS, strongly suggesting that the protomers observed by DLS and EM are the same. Combinations of other charged GFP states can be combined to form similar but less stable protomer particles (**Supplementary Fig. 8**). This is the first example of a defined SUpercharged PRotein Assembly, or SuPrA.

**Fig. 3.**
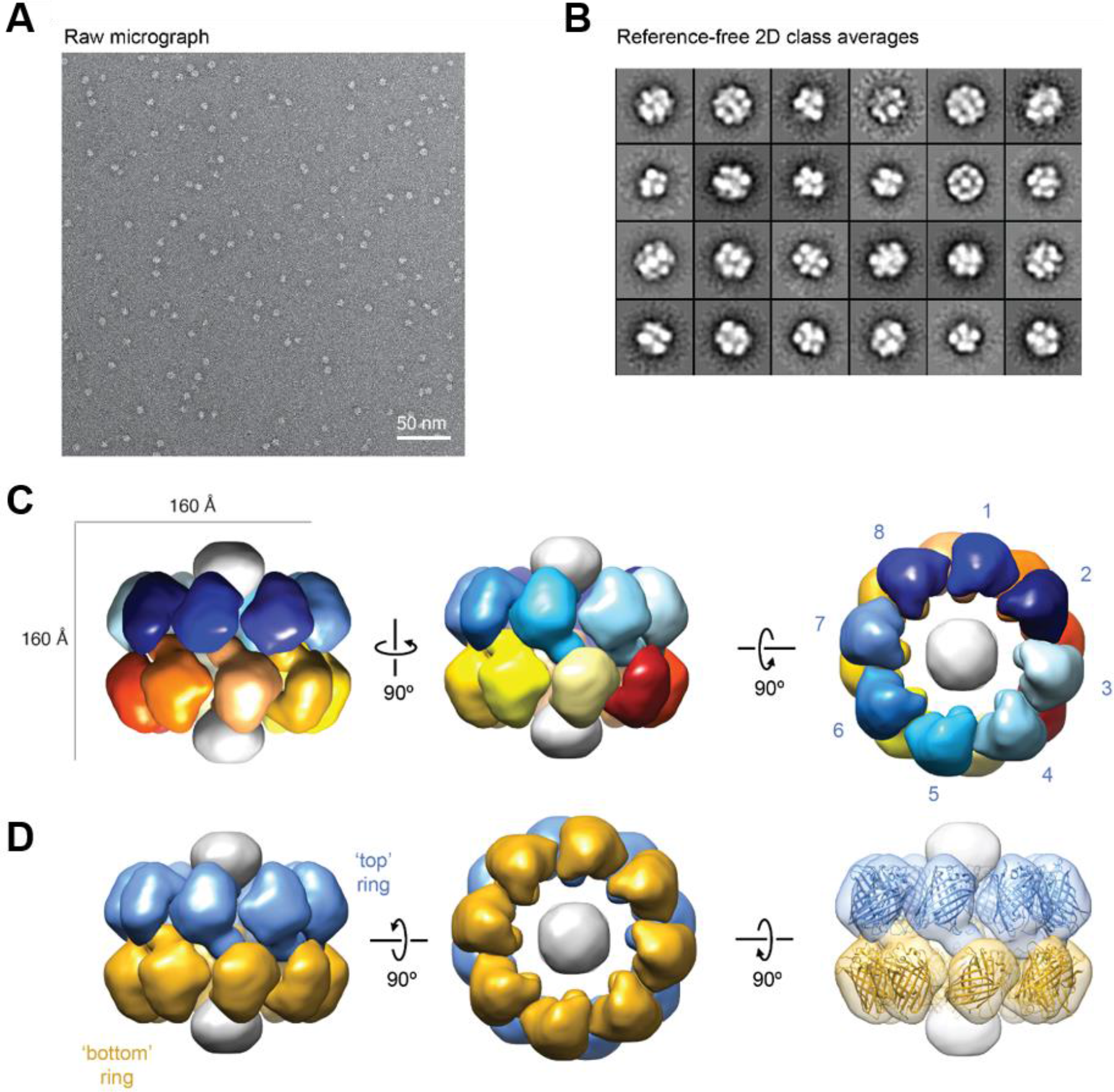
Molecular architecture of the GFP protomer. **a,** Raw micrographs of Ceru+32/GFP-17 particles. **b.,** Reference-free 2D class averages of the protomers **c,** Three-dimensional architecture of negatively stained Ceru+32/GFP-17 protomer at ~18-Å resolution with 8 subunits forming an octameric ring. **d,** Each protomer consists of a top and bottom ring that are offset by ~0.5 subunits in the axial direction. The x-ray crystal structure of GFP (PDB 1EMA) is easily accommodated within the map of each of these subunits.

To further investigate the composition and electronic structures of the SuPrA particles we employed FRET. Exciting Ceru+32/GFP-17 particles at Ceru+32’s absorption maximum, 433 nm, produces GFP-17 emission between 515-535 nm ~30% higher than the sum of the individual spectra, indicating effective energy transfer consistent with separation <10 nm.^50^ To quantify our results, we define a “transfer ratio” of GFP emission in response to 433 nm to that in response to direct excitation (485 nm). Mixed Ceru+32/GFP-17 solutions produces an average transfer ratio of ~0.35, while mixed Ceru+32/GFP+17 solutions produces an average transfer ratio of 0.19, only slightly higher than the ~0.17 value seen for GFP-only solutions (**Supplementary Fig. 9**).

Bulk FRET of Ceru+32/GFP-17 SuPrA particles show a qualitatively similar but more graded dependence on salt concentration compared to DLS. At NaCl ≤50 mM, the transfer ratio remains ~ 0.35 (**Fig. 4a, b**). Increasing NaCl progressively decreases fluorescence transfer until ~450 mM, at which transfer ratios leveled off at ~ 0.22. Notably, increasing NaCl does not affect fluorescence from direct GFP excitation (**Supplementary Fig. 10**). This graded transition suggests that increasing NaCl produces subtle shifts in equilibria or geometries DLS misses. This is not surprising: whereas DLS generally provides the “major” component in a solution via the time-correlation of scattering patterns,^49^ FRET generally provides the “average” component via total energy transfer to acceptor fluorophores. Together, our DLS and FRET data are consistent with subtle but significant shifts in spectral overlap or packing geometries within SuPrA particles of a given size, or with changes in equilibria between differently-sized SuPrA particles.

**Fig. 4.**
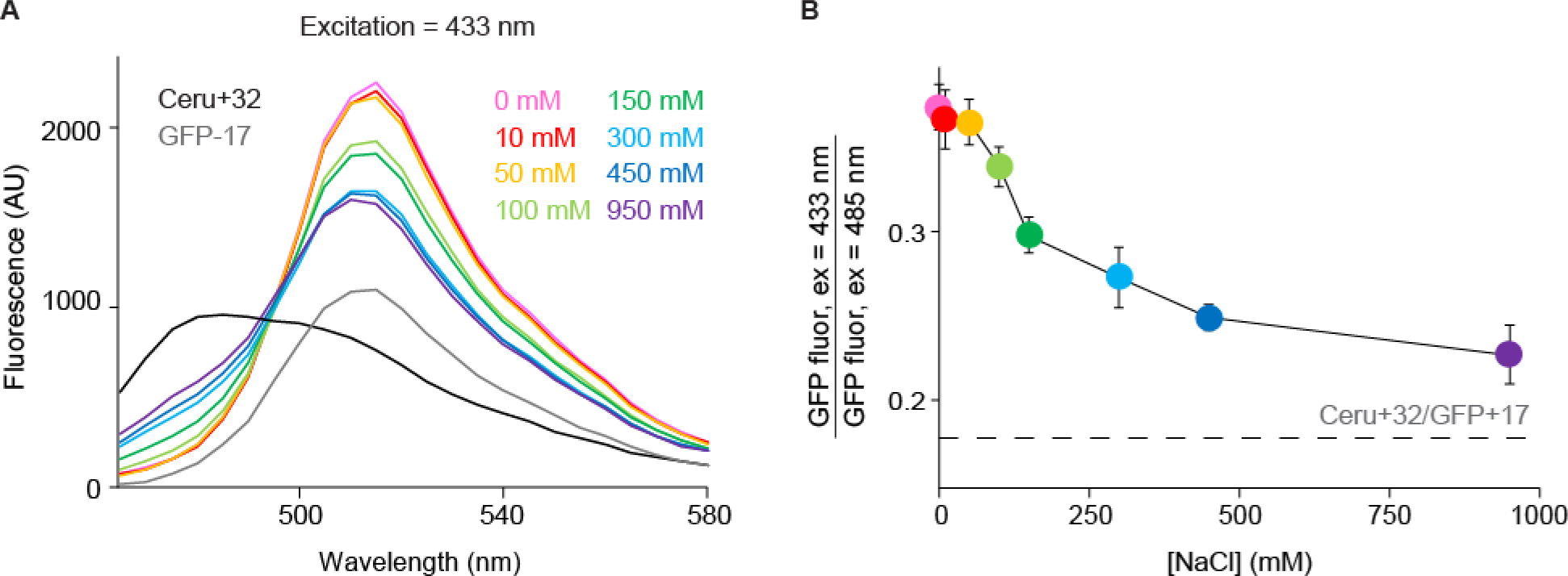
Förster Resonance Energy Transfer (FRET) measurements of Ceru+32/GFP-17 particles show increasing energy transfer between Ceru+32 and GFP-17 with decreasing sodium chloride concentration. **a,** Excitation of Ceru+32/GFP-17 particles at 433 nm, Ceru+32’s excitation maximum, produces decreased Ceru+32 fluorescence and increased GFP-17 fluorescence, indicating transfer. **b,** Increasing salt concentrations progressively decreases fluorescence transfer until around 450 mM where it levels off at approximately 0.22. This graded transition suggests that increasing salt concentrations produce subtle shifts in particle sizes and geometries missed by DLS.

The observation that two oppositely supercharged proteins are required to form SuPrA particles larger than the monomer indicates that assembly is driven by favorable electrostatic interactions. However, the specific interactions driving the formation of a discrete structure rather than a polydisperse variety of shapes and sizes remains unclear. One possibility is that the oligomeric structure arises serendipitously due to specific contacts mediated by introduced residues. An alternative possibility is that while the introduced charged residues provide attractive electrostatics between Ceru+32/GFP-18 interaction, a more general set of shape, patchy steric, or hydrophobic interactions guide formation into a stacked octamer structure. We assessed this hypothesis by assaying the assembly of four “revertant” Ceru+32 variants that each reverted two or three spatially-close residues. We also tested the assembly of a positively supercharged GFP+33 that contained a largely different set of charged residues than did Ceru+32 variant (**Supplementary Fig. 11**).^40^ Upon mixture with GFP-17 at moderate NaCl concentrations each of these variants again forms a ~12 nm structure consistent with the size of the SuPrA protomer, indicating that while charge:charge interactions drive structure formation, the overall shape of the fluorescent protein is sufficient to mediate SuPrA protomer formation. These observations have important implications for further generalizing the methods we describe for nanoparticle formation.

Modeling of SuPrA protomer assembly further indicates that electrostatics and general shape complementarity support the formation of stacked-octamer protomer structures. Specifically, we employed a computational framework developed by the Glotzer lab to robustly assess the assembly of computationally-derived polyhedral shapes to test the stability of simplified candidate protomer structures. Ceru+32 and GFP-17 were modeled as hard, nonconvex polyhedra representing their solvent-excluded surface, with point charges placed at each atomic site (**Fig. 5a**). Because the vast majority of natural homomeric protein complexes contain symmetrical arrangements of proteins,^53^ The total energy at each step was calculated as the sum of the hard particle and screened electrostatic contributions. At the end of the simulation protomer stability was assessed by calculating the radii of gyration *R*_*g*_ (Supp. Eq. 3), defined as the root mean squared molecule-to-molecule distance. The TEM data indicate a *R*_*g*_ ~6.5 nm in the intact SuPrA protomer; we define clear dissociation at *R*_*g*_ > 13 nm. Each candidate configuration remained intact at low NaCl concentrations ([NaCl] < 38mM), again demonstrating that electrostatic and large-scale shape interactions supported stable SuPrA protomer formation. Control SuPrA protomers containing entirely Ceru+32, entirely GFP-17, or an octamer of Ceru+32 juxtaposed with an octamer of GFP-17 were unstable at all NaCl concentrations, consistent with the experimental observations (**Supplementary Fig. 12, 13**) Finer-grained structural modeling (i.e., of hydrophobic effects and counterion release) must await specific structural and physical details that should become manifest with further imaging (i.e., cryo-EM) experiments that better determine the precise symmetric arrangement of monomers.

**Fig. 5.**
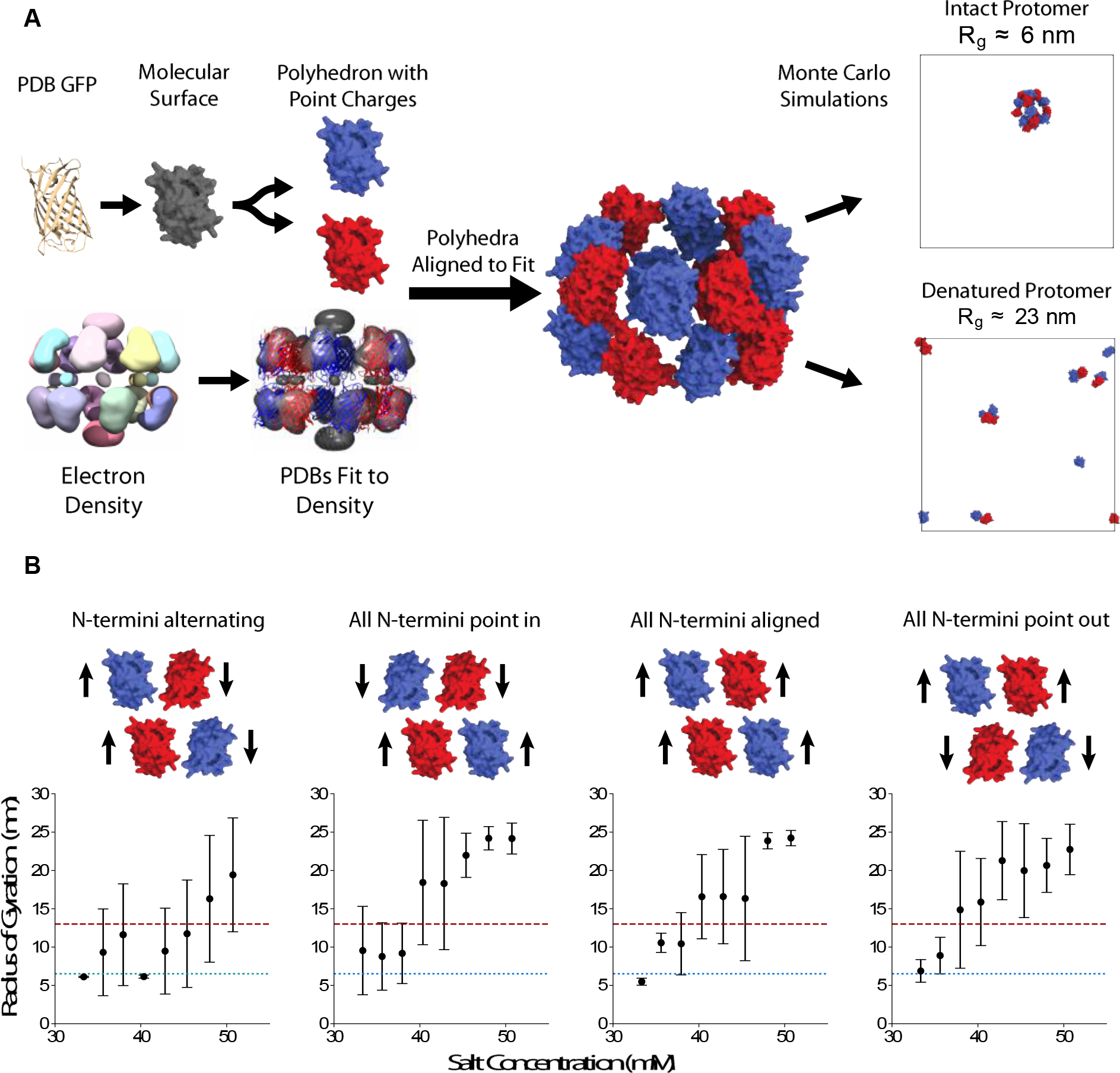
Coarse-grained simulations of candidate protomer structures qualitatively recapitulate experimental data. **a,** To construct coarse-grained protomers, we first generate polyhedral representations of molecular surfaces for the constituent Ceru+32 and GFP-17 structures and assign point charges to each atomic locus. We determine the positions of proteins in the ring by fitting the all-atom representation of the proteins to the electron density. We then use this data to arrange the polyhedra into candidate protomer configurations. We assessed the stability of each configuration by modeling the position of the Ceru+32 and GFP-17 proteins after 400,000 Monte Carlo sweeps, employing both local and geometric cluster moves to equilibrate candidate configurations. **b,** We identified and tested four candidate configurations that all contain alternating, symmetric arrangements of eight Ceru+32 and eight GFP-17 molecules but differ in individual protein orientations. The simulations indicate that all candidate configurations remain intact at low NaCl concentrations, <38 mM but dissolve at higher NaCl concentrations. The highest stable sodium concentration is attained by the conformation in which the orientations of the N-termini alternate within the octameric ring, which dissolves at NaCl ~ 48 mM. Notably, this is still well below the experimentally observed stability limit of ~300 mM.

While previous studies have shown that it is possible to design synthetic protein oligomers either by modifying extant structures^58^.

## Acknowledgements

This material is based upon work supported by the U. S. Army Research Laboratory and the U. S. Army Research Office under grant number W911NF-1-51-0120 to the University of Texas at Austin and under W911NF-15-1-0185 to the University of Michigan. Computational resources and services for simulation work were supported by Advanced Research Computing at the University of Michigan, Ann Arbor. This work used the Extreme Science and Engineering Discovery Environment (XSEDE), which is supported by National Science Foundation grant number ACI-1053575; XSEDE award DMR 140129. AJS is supported by an Arnold O. Beckman Postdoctoral Fellowship. D.W.T is a CPRIT Scholar supported by the Cancer Prevention and Research Institute of Texas (RR160088). This work was supported in part by a Welch Foundation Grant F-1938 (to D.W.T.) The authors thank Barton Dear for helpful discussions regarding interpretation of DLS data and the Texas Materials Institute, part of the Material Science Engineering program at UT-Austin, for supporting the management of the DLS.

## Author Contributions

A.J.S., V.R., J.Gl., A.P., C.J., J.Go., D.W.T., S.C.G, and A.D.E. conceived and designed the experiments. A.P., A.J.S. and J.Go. designed proteins. A.J.S., B.R.M and A.P. expressed proteins. A.P., J.Go., and C.J. performed early optimization of DLS and FRET experiments. A.J.S. carried out DLS and FRET experiments and analyzed the data. J.Ge, J.L., and D.W.T. performed EM experiments and analyzed data. V.R. and J.Gl. designed the simulations. V.R. performed simulations. A.J.S., V.R., J.Gl., S.C.G, and A.D.E. wrote the manuscript, and all authors reviewed and commented upon the manuscript.

## References

[1] Whitesides, G.M. & Grzybowski, B. Self-assembly at all scales. Science 295, 2418–2421 (2002).

[2] Blundell, T. L. & Srinivasan, N. Symmetry, stability, and dynamics of multidomain and multicomponent protein systems. Proc Natl Acad Sci USA 93, 14243–14248 (1996).

[3] Levy, E. D., Boeri Erba, E., Robinson, C.V., Teichmann, S. A. Assembly reflects evolution of protein complexes. Nature 453, 1262–1265 (2008).

[4] Goodsell, D. S. & Olson, A. J. Structural symmetry and protein function. Annu Rev Biophys Biomol Struct 29, 105–153 (2000).

[5] André, I., Strauss, C. E. M., Kaplan, D. B., Bradley, P. & Baker, D. Emergence of symmetry in homooligomeric biological assemblies. Proc. Natl. Acad. Sci. 105, 16148–16152 (2008).

[6] Plaxco, K. W. & Gross, M. Protein Complexes: The Evolution of Symmetry. Curr. Biol. 19, R25–R26 (2009).

[7] Bergendahl, L. T., & Marsh, J. H. Functional determinants of protein assembly into homomeric components. Sci Rep 7, 4932 (2017).

[8] Padilla, J.E., Colovos, C., & Yeates, T.O. Nanohedra: using symmetry to design self assembling protein cages, layers, crystals, and filaments. Proc Natl Acad Sci USA 98, 2217–2221 (2001).

[9] Lai, Y-T., Cascio, D., & Yeates, T.O. Structure of a 16-nm cage designed by using protein oligomers. Science 336, 1129 (2012)

[10] Bale, J. B. et al. Accurate design of megadalton-scale two-component icosahedral protein complexes. Science 353, 389–394 (2016)

[11] Hsia, Y. et al. Design of a hyperstable 60-subunit protein icosahedron. Nature 535, 136–139 (2016)

[12] Butterfield, G.L., et al. Evolution of a designed protein assembly encapsulating its own RNA genome. Nature 552, 415–420 (2017).

[13] Alberstein, R., Suzuki, Y., Paesani, F., & Tezcan, F. A. Engineering the entropy-driven free-energy landscape of a dynamic nanoporous protein assembly. Nat Chem 10.1038/s41557-018-0053-4 (2018).

[14] Badieyan, S., et al. Symmetry - directed self - assembly of a tetrahedral protein cage mediated by de novo - designed coiled coils. ChemBioChem 18, 1888–1892 (2017).

[15] Brodin, J.D., Carr, J.R., Sontz, P.A., & F. A. Tezcan. Exceptionally stable, redox-active supramolecular protein assemblies with emergent properties. Proc Nat Acad Sci USA 111, 2897–2902 (2014)

[16] Butterfield, G.L., et al. Evolution of a designed protein assembly encapsulating its own RNA genome. Nature 552, 415–420 (2017).

[17] Yeates, T.O. Geometric principles for designing highly symmetric self-assembling protein nanomaterials. Ann. Rev. Biophys. 46:23–42 (2017). Kobayashi, N. & Arai, R. Design and construction of self-assembling supramolecular protein complexes using artificial and fusion proteins as nanoscale building blocks. Curr Op Biotech 46, 57–65 (2017).

[137,18] Damasceno, P.F., Engel, M., & Glotzer, S.C. Predictive self-assembly of polyhedral into complex structures. Science 337, 453–457 (2012).

[19] Paik, T. & Murray, C. B. Shape directed binary assembly of anisotropic nanoplates: A nanocrystal puzzle with shape-complementary building blocks. Nano Lett 13, 2952–2956 (2013).

[20] Gong, J. et al. Shape-dependent ordering of gold nanocrystals into large-scale superlattices. Nat Comm 8, 14038 (2017).

[21] Wolters, J. R., et al. Self-assembly of “Mickey Mouse” shaped colloids into tube-like structures: experiments and simulations. Soft Matter 11, 1067–1077 (2015).

[22] Boles, M.A., Engel, M., & Talapin, D.V. Self-assembly of colloidal nanocrystals: From intricate structures to functional materials. Chem Rev 116, 11220–11289 (2016).

[23] Fu, L. et al. Assembly of hard spheres in a cylinder: a computational and experimental study. Soft Matter 13, 3296–3306 (2017).

[24] Ye, X., et al. Competition of shape and interaction patchiness for self-assembling nanoplates. Nat Chem 5, 466–473 (2013).

[25] Zhang, Z. & Glotzer, S.C. Self-assembly of patchy particles. Nano Lett. 4, 1407–1413 (2004).

[26] Giacometti, A., Lado, F., Largo, J., Pastore, G. & Sciortino, F. Effects of patch size and number within a simple model of patchy colloids. J. Chem. Phys. 132, 174110 (2010).

[27] Pawar, A.B. & Kretzschmar, I. Fabrication, Assembly, and Application of Patchy Particles. Macromol. Rapid Comm. 31, 150–168 (2010).

[28] Gong, Z., Hueckel, T., Yi, G.R. & Sacanna, S. Patchy particles made by colloidal fusion. Nature 550, 234–238 (2017).

[29] Duguet, E., Hubert, C., Chomette, C., Perroc, A. & Ravaine, S. Patchy colloidal particles for programmed self-assembly. Comptes Rendus Chimie 19, 172–182 (2016).

[30] Woo, S., & Rothemund, P.K. Programmable molecular recognition based on the geometry of DNA nanostructures. Nat Chem 3, 620–627 (2011).

[31] Gerling, T., Wagenbauer, K.F., Neuner, A.M., Dietz, H. Dynamic DNA devices and assemblies formed by shape-complementary, non-base pairing 3D components. Science 347, 1446–1452 (2015).

[32] Sheinerman, F.B., Norel, R., & Honig, B. Electrostatic aspects of protein-protein interactions. Curr Op Struct Biol 10, 153–159 (2000).

[33] Liljeström, V., Mikkilä, J., & Kostiainen, M.A. Self-assembly and modular functionalization of threedimensional crystals from oppositely charged proteins. Nat Commun 5, 4445 (2014).

[34] Kostiainen, M.A et al. Electrostatic assembly of binary nanoparticle superlattices using protein cages. Nat Nanotechnol 8, 52–56 (2012).

[35] Seebeck, F. P. et al. A simple tagging system for protein encapsulation. J Am Chem Soc 128, 4516–4517 (2006).

[36] Wörsdörfer, B., Woycechowsky, K. J. & Hilvert, D. Directed evolution of a protein container. Science 331, 589–592 (2011).

[37] Held, M., Kolb, A., Perdue, S., Hsu, S.-Y., Bloch, S.E., Quin, M.B., & Schmidt-Danner, C. Engineering formation of multiple recombinant Eut protein nanocompartments in E. coli. Sci Rep 6, 24359 (2016).

[38] Beck, T., Tetter, S., Künzle, M., & Hilvert, D. Construction of Matryoshka-type structures from supercharged protein nanocages. Angew Chemie 54, 937–940 (2014).

[39] Sun, H., Luo, Q., Hou, C. & Liu, J. Nanostructures based on protein self-assembly: From hierarchical construction to bioinspired materials. Nanotoday 14, 16–41 (2017).

[40] Lawrence, M.S., Phillips, K.J. & Liu, D.R. Supercharging proteins can impart unusual resilience. J. Am. Chem. Soc. 129, 10110–10112 (2007).

[41] Der, B.S. et al. Alternative computational protocols for supercharging protein surfaces for reversible unfolding and retention of stability. PLoS One 31, e64363 (2013).

[42] Oh, H.J., Gather, M.C., Song, J.-J. & Yun, S.H. Lasing from fluorescent protein crystals. Opt. Express, 22, 31411–31416 (2014).

[43] Ormö, M. et al. Crystal structure of the Aequorea victoria green fluorescent protein. Science 273, 1392–1395 (1996).

[44] Bolhuis, P. & Frenkel, D. Tracing the phase boundaries of hard spherocylinders. J. Chem. Phys. 106, 666–687.

[45] Rizzo, M.A., Springer, G.H., Granada, B. & Piston, D.W. An improved cyan fluorescent protein variant useful for FRET. Nat. Biotech. 4, 445–449 (2004).

[46] Goedart, J. et al. Structure-guided evolution of cyan fluorescent proteins towards a quantum yield of 93%. Nat. Commun. 3, 751 (2012).

[47] Goedhart, J. et al. Bright cyan fluorescent protein variants identified by fluorescence lifetime screening. Nat. Met. 7, 137–139 (2010).

[48] García-Seisdedos, H., Empereur-Mot, C., Elad, N. & Levy, E.D. Proteins evolve on the edge of supramolecular self-assemble. Nature 548, 244–247 (2017).

[49] Hassan, P.A., Rana, S. & Verma, G. Making sense of Brownian motion: colloid characterization by dynamic light scattering. Langmuir 31, 3–12 (2015).

[50] Bajar, B.T., Wang, E.S., Zhang, S., Lin, M.Z. & Chu, J. A guide to fluorescent protein FRET pairs. Sensors 16, 1488 (2016).

[51] Anderson, J. A., Lorenz, C. D., & Travesset, A. General purpose molecular dynamics simu-lations fully implemented on graphics processing units. J. Comp. Phys 227, 5342–5359 (2008).

[52] Glaser, J., Nguyen T. D., Anderson, J. A., Liu, P., Spiga, F., Millan, J.A., Morse, D. C., & Glotzer, S.Cs. Strong scaling of general-purpose molecular dynamics simulations on GPUs. Comp Phys Commun 192, 97–107 (2015).

[53] Sinkovits, D.W., Barr, S.A., & Luijten, E. Rejection-free Monte Carlo scheme for anisotropic particles. J Chem Phys 136, 144111 (2012).

[54] Yang, F., Moss, L.G., & Phillips Jr., G.B. The molecular structure of green fluorescent protein. Nat Biotechnol 14, 1246–1251 (1996)/

[56] Lyskov, S. et al. Serverification of molecular modeling applications: The Rosetta online server that includes everyone (ROSIE). PLoS One 8, e63906 (2013).

[57] Miklos, A.E. et al. Structure-based design of supercharged, highly thermoresistant antibodies. Chem Biochem 19, 449–455 (2012).

[58] Johnson, L.B., Park, S., Gintner, L.P., & Snow, C. D. Characterization of supercharged cellulase activity and stability in ionic liquids. Journal of Molecular Catalysis B: Enzymatic 132, 84–90 (2016).

